# Structural Modeling Identifies a Conserved ADAR1 dsRBDIII Motif Predicted to Interact with PKR at Thr451

**DOI:** 10.64898/2026.10.23.761487

**Authors:** Saman Taj, Meng Hao, Antonija Braut, Igor Jurak

## Abstract

Protein kinase R (PKR) is a key cellular dsRNA sensor whose activation is regulated by ADAR1. Although ADAR1–PKR association is established, its molecular determinants remain poorly understood. Molecular docking identified a conserved W768/F769/P770 (WFP) motif within ADAR1 dsRBD III as a potential PKR-interaction interface. Disruption of this motif altered the predicted binding configuration. However, co-immunoprecipitation revealed no significant reduction in PKR association following individual substitutions or deletion of the WFP motif, although the deletion showed a consistent trend toward reduced binding. RNase treatment reduced PKR co-immunoprecipitation approximately tenfold without abolishing the interaction. These findings suggest that ADAR1–PKR association is predominantly RNA-dependent, potentially involving shared dsRNA substrates that facilitate direct protein–protein contacts, with the WFP motif contributing to interaction stability.

Graphical Abstract:
Schematic representation of ADAR1p150 and PKR domain organization, predicted ADAR1– PKR interaction interface, and experimental validation.
(Top left) Domain organization of adenosine deaminase acting on RNA 1 p150 (ADAR1p150), showing the N-terminal Z-DNA-binding domains (Zα and Zβ), three double-stranded RNA-binding domains (RI, RII, and RIII), and the C-terminal deaminase domain (DD). The W768/F769/P770 motif (WFP motif) is highlighted in RIII. (Top right) Domain organization of protein kinase R (PKR), showing its two N-terminal double-stranded RNA-binding motifs (RI and RII) and C-terminal kinase domain (KD). The adjacent structural representation highlights the PKR kinase domain and the region containing Thr451 (T451), a residue implicated in PKR activation. (Bottom left) Predicted ADAR1–PKR interaction interface generated by protein–protein docking. ADAR1 RBDIII is shown in green, and the PKR kinase domain is shown in cyan and magenta to distinguish the displayed protein regions or chains. The ADAR1 residues Trp768 (W768), Phe769 (F769), and Pro770 (P770) are highlighted near PKR Thr451, illustrating their proximity within the predicted interface. The RNA strand shown beneath the complex illustrates the context of double-stranded RNA recognition and is not necessarily part of the docked protein–protein complex. (Bottom right) In-vitro co-immunoprecipitation (Co-IP) analysis comparing PKR association with wild-type ADAR1p150 (ADAR1-GFP WT) and the combined WFP mutant (WFP). The bar graph shows the relative PKR signal recovered in ADAR1 immunoprecipitates; error bars indicate variability between measurements, and *ns* denotes a statistically non-significant difference between the groups.

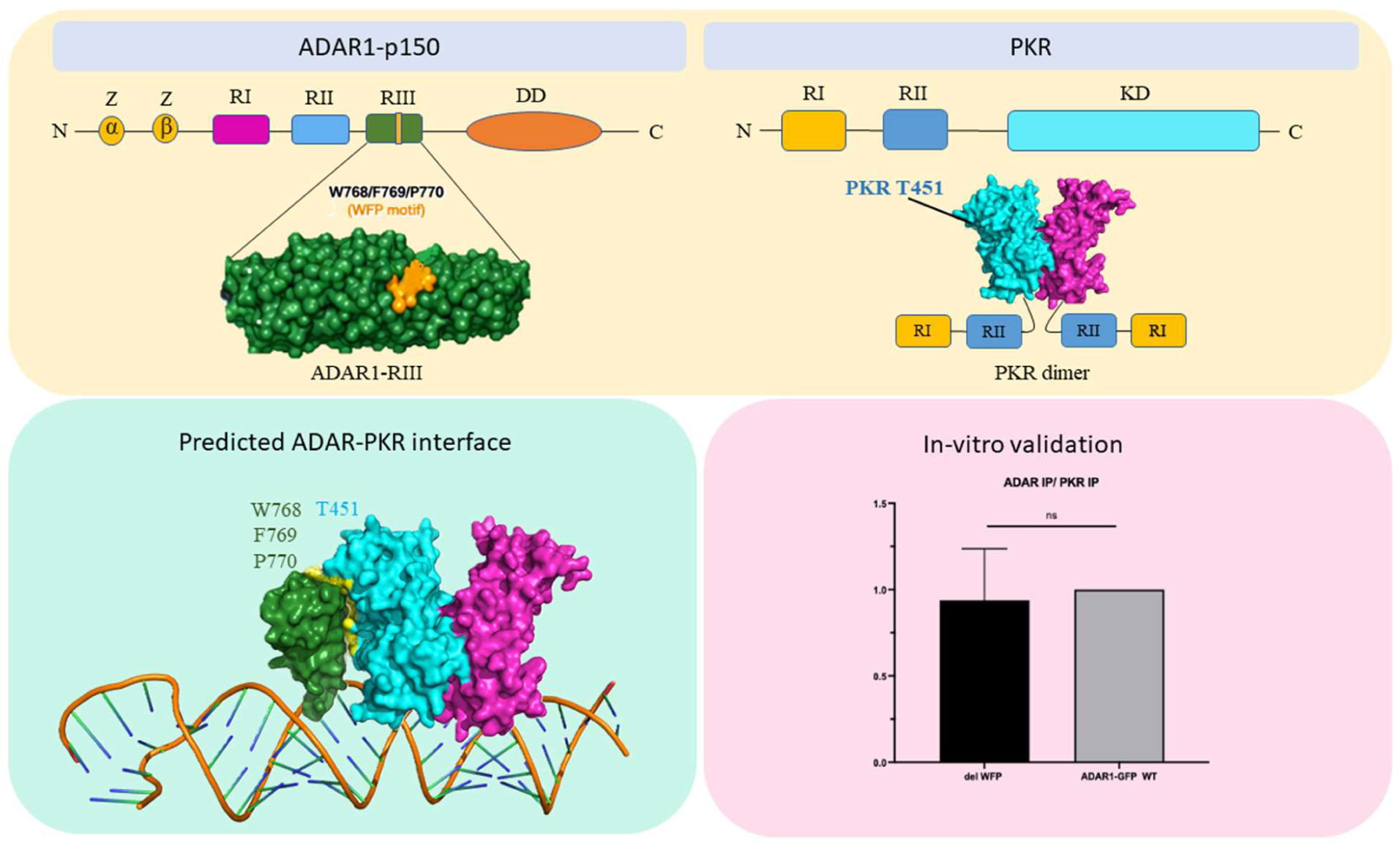

## Introduction

Host-virus interactions have co-evolved over millions of years, resulting in sophisticated cellular mechanisms that detect viral infection and restrict viral replication. Host cells recognize pathogen-associated molecular patterns (PAMPs), including viral nucleic acids, which activate interferon (IFN) responses and establish an antiviral state or, under certain conditions, under certain conditions, induce cell arrest or programmed cell death [1, 2]. Among these molecular patterns, viral double-stranded RNA (dsRNA) is an important trigger of innate antiviral immunity and can arise during viral replication [3]. Multiple host sensors recognize viral dsRNA, including Toll-like receptors (TLRs), RIG-I-like receptors such as retinoic acid-inducible gene I (RIG-I) and melanoma differentiation-associated protein 5 (MDA5), laboratory of genetics and physiology 2 (LGP2), Z-DNA-binding protein 1 (ZBP1), protein kinase R (PKR), and members of the oligoadenylate synthetase (OAS) family [1, 4]. Activation of these sensors initiates signaling pathways that restrict viral replication. However, endogenous cellular dsRNAs can also engage these sensing pathways and induce inappropriate immune activation [5]. To maintain discrimination between self and non-self RNA, cells employ regulatory mechanisms including adenosine-to-inosine (A-to-I) RNA editing catalyzed by adenosine deaminases acting on RNA (ADARs) (reviewed in [6, 7]. Protein kinase R (PKR) is an interferon-induced serine/threonine kinase that contributes to antiviral defense by sensing dsRNA generated during viral infection (reviewed in [8, 9]). PKR contains an N-terminal regulatory region comprising two tandem dsRNA-binding motifs, with motif 1 spanning residues 6–79 and motif 2 spanning residues 96– 169, separated by an inter-motif spacer (IMS) region (Figure 1). Its C-terminal region contains the kinase catalytic domain encompassing residues 274–551 [10]. Following activation, the kinase domain phosphorylates the α-subunit of eukaryotic translation initiation factor 2 (eIF2α), resulting in suppression of translation initiation and thereby limiting viral protein synthesis and replication [11]. In its inactive state, intramolecular interactions involving the regulatory and kinase regions contribute to PKR autoinhibition [12]. Upon binding to sufficiently long dsRNA, PKR undergoes conformational rearrangement and dimerization, facilitating activation of the C-terminal kinase domain. dsRNA molecules of approximately ≥60 bp are generally more effective at promoting PKR activation, whereas shorter or single-stranded RNAs exhibit reduced activity [10, 13–16]. Structural studies have further shown that activation reorganizes the PKR dimer to create a configuration capable of accommodating eIF2α [17, 18], while an specific region within the kinase domain has been implicated in interaction with ADAR1[19]. PKR contains several phosphorylation sites, including S242 within the sequence KAKRSLAPR, T255 within DMKETKYTV, T258 within ETKYTVDKR, T446 within DGKRTRSKG, and T451 within RSKGTLRYM (reviewed in[20]). Among these residues, Thr446 and Thr451 are located within the activation loop of the kinase domain and contribute to PKR activation. Autophosphorylation within this region is required for efficient kinase activation, with Thr446 identified as a major phosphorylation site that contributes to stabilization of the active kinase conformation and substrate binding [18]. Thr451 also contributes to efficient PKR activation and kinase activity [20]. Together, these structural and phosphorylation-dependent changes convert PKR from an inactive sensor into an active kinase capable of phosphorylating eIF2α.

**Figure 1:**
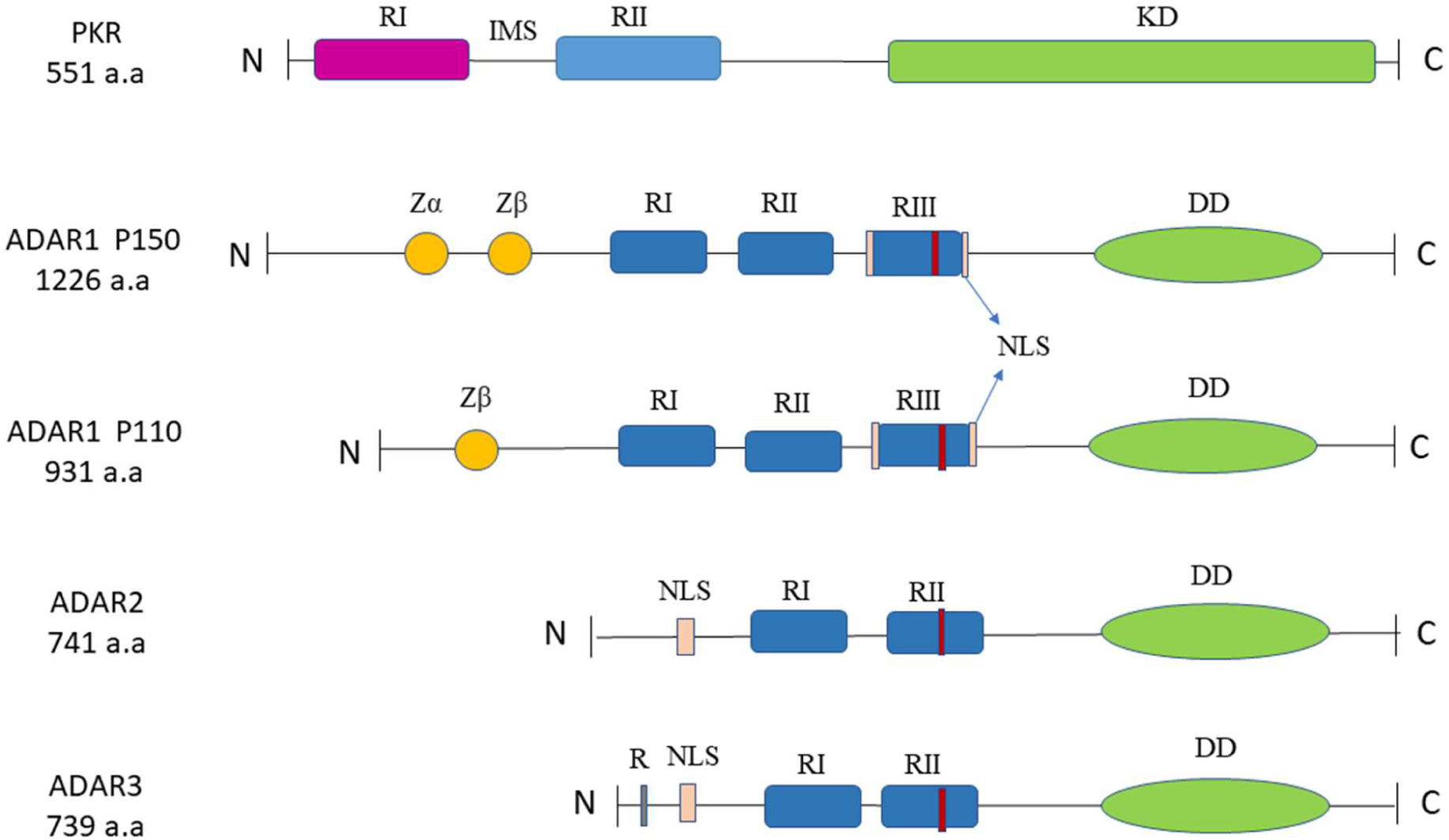
Schematic representation of domain organization in different ADAR proteins and PKR. RI, RII, and RIII indicate RNA-binding domains; Zα and Zβ represent Z-DNA binding domains; DD denotes the deaminase domain; KD represents kinase domain; R marks the site for single- and double-stranded RNA binding; NLS indicates the nuclear localization signal and IMS shows intra motif sequences.

The adenosine deaminase acting on RNA (ADAR) family in mammals comprises ADAR1, ADAR2, and ADAR3, which participate in the recognition and modification of cellular dsRNA [21, 22] ADAR-mediated A-to-I editing can alter RNA structure, stability, and function and is an important mechanism for regulating cellular responses to endogenous dsRNA. ADAR1 exists primarily as two isoforms, p150 and p110, which differ in their N-terminal regions. ADAR1p150 contains two Z-nucleic-acid-binding domains, Zα and Zβ, whereas ADAR1p110 lacks the N-terminal Zα domain. Both isoforms contain three dsRNA-binding domains, designated RBDI, RBDII, and RBDIII, followed by a C-terminal deaminase domain responsible for catalytic RNA editing. The third RNA-binding domain of ADAR1 encompasses residues A742–G794 and is positioned immediately upstream of the linker connecting the RNA-binding region to the catalytic domain [23] (Figure 1). In contrast, ADAR2 and ADAR3 contain two dsRNA-binding domains and lack the N-terminal Zα/Zβ domain arrangement characteristic of ADAR1 [22] (Figure2). A substantial body of evidence demonstrates that ADAR1p150 exerts functions beyond its catalytic RNA-editing activity. ADAR1 can regulate several cytosolic dsRNA-sensing pathways, including those mediated by MDA5, RIG-I, ZBP1, OAS proteins, and PKR, through mechanisms that are not necessarily dependent on A-to-I editing [24–27]. Among the dsRNA-sensing pathways regulated by ADAR1, the ADAR1–PKR interaction is of particular interest because it can limit excessive PKR activation and its associated translational suppression [28]. Recent structural and biochemical evidence provides direct support for an ADAR1-PKR interaction[19, 24, 28]. The third RNA-binding domain of ADAR1 (RBDIII) was shown to interact with the kinase domain of PKR, and disruption of this interaction impaired ADAR1-mediated suppression of PKR activation [19]. These findings identify ADAR1 RBDIII as an important regulatory region and support a model in which direct molecular contacts between ADAR1 and PKR contribute to modulation of kinase activity. However, although the interaction has been demonstrated experimentally, the molecular determinants governing the ADAR1-PKR interface remain incompletely characterized. In particular, the specific residues within ADAR1 RBDIII and the PKR kinase domain that contribute to the interaction, the nature of their molecular contacts, and their contribution to complex stability and PKR regulation require further investigation. Defining these determinants is particularly important because the ADAR1-PKR interaction occurs in the context of dsRNA recognition, making it necessary to distinguish direct protein-protein contacts from effects arising from RNA binding or RNA editing.

In the present study, we investigated the molecular basis of the interaction between ADAR1 RBDIII and the PKR kinase domain using protein-protein docking. A structural model of the complex was generated to identify key interfacial residues and characterize their predicted molecular contacts. Based on the docking model, selected ADAR1 residues were targeted for mutational analysis to evaluate their contribution to ADAR1-PKR association. By integrating structural modelling with experimental analysis of protein association, this study identifies candidate molecular determinants of the ADAR1-PKR interface and provides mechanistic insight into the editing-independent regulation of PKR by ADAR1.

## Materials and Methods

### Protein Structure Retrieval and Preparation

The three-dimensional (3D) structures of ADAR1 (PDB ID: 7ZLQ) and PKR (PDB ID: 2A19) were retrieved from the Protein Data Bank (PDB). Prior to docking, structural optimization was performed using the UCSF Chimera (version 3.31). Bound ligands, solvent molecules, and other non-protein components were removed prior to docking. The PKR structure comprises three chains; Chain A, corresponding to the bound eIF2α substrate, was excluded, whereas Chains B and C, representing the catalytically active kinase-domain dimer, were retained for docking (Dar et al., 2005). The dsRNA molecule bound to the RBDIII domain of ADAR1 was also removed to permit protein-protein docking. Both protein structures were protonated and energy-minimized using the MMFF94x force field with partial atomic charges. Energy minimization was performed using the steepest descent algorithm followed by the conjugate gradient algorithm until a convergence criterion of 0.1 kcal/mol/Å was reached[29, 30].

### Molecular Docking

Protein–protein docking was performed using the HDOCK server (http://hdock.phys.hust.edu.cn/), which utilizes a hybrid docking algorithm that integrates template-based modeling with free docking strategies to predict protein–protein interactions. The optimized protein structures were submitted in PDB format, with PKR designated as the receptor and ADAR1 as the ligand and Protein–protein docking was performed using the default parameters. The HDOCK server generated multiple docking conformations that were ranked according to docking scores, which reflect predicted binding affinity and interaction stability. Docking models were evaluated based on docking score, confidence score, and predicted interaction interface. Protein–protein interfaces, interacting residues, and inter-residue distances were analyzed and visualized using UCSF Chimera (version 3.31) and PyMOL (V4.6.0).

### Sequence Alignment and Evolutionary Conservation Analysis

To investigate sequence conservation of the predicted PKR-interaction motif, protein sequences of human ADAR family members, including ADAR1 (UniProt: P55265), ADAR2 (UniProt: P78563), and ADAR3 (UniProt: Q9NS39), were retrieved from the UniProt database (https://www.uniprot.org/) [21]. To evaluate evolutionary conservation, ADAR1 orthologs were additionally obtained from human (*Homo sapiens*; UniProt: P55265), chimpanzee (*Pan troglodytes*; UniProt: A0A2I3T9M4), mouse (*Mus musculus*; UniProt: Q99MU3), chicken (*Gallus gallus*; UniProt: A0A1D6Y7R2), and zebrafish (*Danio rerio*; UniProt: A0A8M1P8J0). To assess conservation of the motif in the absence of PKR, ADAR homologs from representative metazoan species lacking canonical PKR orthologs, including fruit fly (*Drosophila melanogaster*; UniProt: Q9NII1), nematode (*Caenorhabditis elegans*; UniProt: Q9U3D6), and hydra (*Hydra vulgaris*; UniProt: T2M593), were also included in the analysis. Multiple sequence alignment of all protein sequences was performed using the Clustal Omega online server (https://www.ebi.ac.uk/jdispatcher/msa/clustalo) with default parameters. The resulting alignments were analyzed to identify conserved residues and motifs within the dsRNA-binding domains, with particular emphasis on the predicted ADAR1–PKR interaction interface.

### Comparative Docking of ADAR Family Members

To compare the predicted interaction of ADAR family members with PKR, protein–protein docking analyses were performed for ADAR2 and ADAR3 using the same computational workflow applied to ADAR1. The crystal structure of the human ADAR2 dsRNA-binding domain was obtained from the Protein Data Bank (PDB ID: 8E0F). As no experimentally determined three-dimensional structure of human ADAR3 is available in the Protein Data Bank (PDB), the predicted structure of ADAR3 (UniProt: Q9NS39; AlphaFold model: AF-Q9NS39-F1) was retrieved from the AlphaFold Protein Structure Database (Jumper et al., 2021; Bertoni et al., 2025). The previously prepared PKR kinase-domain dimer (PDB ID: 2A19; Chains B and C) was used as the receptor for all comparative docking analyses. Prior to docking, The ADAR2 structure was prepared following the same protocol described for ADAR1. The AlphaFold-predicted ADAR3 structure did not contain bound ligands or solvent molecules and was therefore used directly. Protein–protein docking of ADAR2 and ADAR3 with PKR was performed using the HDOCK server following the same protocol described above. PKR was designated as the receptor, whereas ADAR2 and ADAR3 were used as ligands in separate docking simulations. Multiple docking conformations were generated and ranked according to docking score. The highest-ranked docking model for each complex was selected for subsequent structural comparison and interaction analysis.

### Site-Directed Mutagenesis and Re-docking

To investigate the functional significance of the conserved motif GRWFPA within the RBDIII domain of ADAR1, site-directed mutagenesis was performed using the PyMOL Mutagenesis Wizard. Three conserved residues within the GRWFPA motif were simultaneously substituted to generate the sequence GR<u>EYG</u>A, where selected residues were altered to disrupt potential on interfaces. The substitutions comprised W768E, F769Y, and P770G, and the modified ADAR1 RBDIII structure was saved in PDB format for subsequent docking analysis. The mutated ADAR1 RBDIII structure was subsequently subjected to protein–protein docking with PKR (chain B and C) using the HDOCK server (http://hdock.phys.hust.edu.cn/), under default parameters. The resulting docking complexes were analyzed to compare the predicted interaction interface with that of the wild-type complex.

### Cells and plasmids

ADAR1 KO cells (derivatives of HEK-293 cells [28, 31] were cultured in Dulbecco’s Modified Eagle Medium (PAN Biotech) supplemented with 10% fetal bovine serum (PAN Biotech), 100 µg/mL Penicillin-Streptomycin (Capricorn), 2 mM L-Glutamine (Capricorn), and 1 mM Sodium Pyruvate (Capricorn) under standard culture conditions in a humidified incubator at 37°C and 5% CO2. The expression vector pmGFP-ADAR1-p150 (Addgene #117927) encoding the human ADAR1 p150 isoform was constructed by the Kumiko Ui-Tei Laboratory. Point mutants (W768E, F769Y, P770G, K778E, and R790E) and the deletion mutant (ΔWFP) were generated from the pmGFP-ADAR1-p150 backbone by GenScrip. An empty GFP vector (pEGFP-N1) was included as a negative control.

### Plasmid Transfection

ADAR1 KO cells were seeded in 6-well plates 24 hours prior to transfection. Transfection with the indicated plasmid constructs (pmGFP control, wild-type pmGFP-ADAR1-p150, or ADAR1- p150 mutant plasmids) was performed using Lipofectamine 2000 (Invitrogen #11668-027) according to the manufacturer’s instructions.

### Co-Immunoprecipitation (Co-IP)

For protein-protein interaction analysis between PKR and GFP-ADAR1 variants, transfected ADAR1 KO cells were lysed in RIPA buffer (50 mM Tris pH 8.0, 150 mM NaCl, 1% NP-40, 0.5% sodium deoxycholate, 0.1% SDS) supplemented with complete protease inhibitor cocktail (Roche). Cell lysates were cleared by centrifugation at 14000 rpm for 10 minutes at 4°C. The supernatant of each sample was separated into two fractions. 15 uL was frozen as a total protein fraction while the remaining 85uL was mixed with 3uL GFP antibody (Santa Cruz, #sc-9996) and kept on gentle rolling overnight at 4°C. The next day, 50 µL of Dynabeads Protein G (Invitrogen #10004D) was added to the mixture were incubated on roller on RT for at least another 4h before washing. The magnetic beads were washed five times with RIPA buffer and boiled at 95°C for 7 minutes in 4× Laemmli sample buffer (Bio-Rad) to elute bound protein complexes for analysis. Primary antibodies used for immunoprecipitation and Western blot included rabbit anti-PKR (Cell Signaling Technology, #12297S), rabbit anti-ADAR1 (Cell Signaling Technology, #32136S for p150), and mouse anti-GFP (Santa Cruz Biotechnology, #sc-9996). Horseradish peroxidase (HRP)-conjugated goat anti-rabbit IgG (#7074S) and horse anti-mouse IgG (#7076S; 1:2,000) from Cell Signaling Technology were used as secondary antibodies.

### Western blot

For Western blot analysis cells were lysed in RIPA buffer (150mM NaCl, 1% NP-40, 0.5% Na deoxycholate, 0.1% SDS, 50mM Tris (pH 8.0)) supplemented with Complete protease inhibitor (Roche) and mixed with 4x Laemelli buffer. The proteins were separated in 10% denaturing PAGE and transferred to a nitrocellulose membrane (Machery-Nagel) and Western blotting was performed as previously described [32]. Proteins were visualized with Amersham ECL (Cytiva) or Supersignal West Femto Detection Reagent (Thermo Fischer Sc.) in Chemidoc (BioRAD).

## Results

### ADAR1 and PKR interaction

The regulation of PKR by ADAR1 is essential for the control of innate immunity and the maintenance of cellular homeostasis, and their direct protein-protein interaction has been demonstrated in multiple studies [24, 33]. Recently, Sinigaglia et al. used AlphaFold-Multimer (AFM) to predict that the dsRBDIII domain of ADAR1 interacts with the kinase domain of PKR near its activation loop, in a manner analogous to PKR’s intramolecular regulatory interactions [19]. Consistent with this model, ADAR1 dsRBDI and dsRBDII were not predicted to interact with PKR in the absence of dsRBDIII, suggesting a specific role for dsRBDIII in mediating this interaction. Although biochemical studies have demonstrated that ADAR1 antagonizes PKR activity [24, 28, 33], the structural basis of this inhibition remains unclear.

To further characterize the interaction interface between ADAR1 and PKR, we performed targeted protein–protein docking analysis of the PKR and the ADAR1 dsRBDIII domain using the HDOCK online server. Briefly, Molecular docking was performed using the prepared three-dimensional structures of ADAR1 (PDB ID: 7ZLQ) and PKR (PDB ID: 2A19). The PKR kinase-domain dimer (Chains B and C) was used as the receptor, whereas ADAR1 was used as the ligand [20]. Docking was carried out using the HDOCK server (http://hdock.phys.hust.edu.cn/) following standard structure preparation procedures [34]. The top-ranked docking conformation was selected based on docking score, and interaction pose for subsequent structural and interaction analyses. Surprisingly, in contrast to the AlphaFold-Multimer model proposed by Sinigaglia et al [19], which predicted that ADAR1 dsRBDIII interacts with the PKR kinase domain adjacent to the activation loop and experimentally validated the importance of the ADAR1 interface residues Lys778 and Arg790 for inhibition of PKR autophosphorylation at Thr446, our HDOCK-based protein-protein docking analysis predicted a distinct binding interface. Specifically, the interaction was localized primarily to the cleft between the N- and C-terminal lobes of the PKR kinase domain and involved residues surrounding Thr451 rather than Thr446 (Figure 2). Specifically, the docking model predicted that Thr451 of PKR forms contacts with Trp768, Phe769, Pro770, and Val791 within the dsRBDIII domain of ADAR1. The corresponding inter-residue distances were 4.904 Å, 4.615 Å, 3.991 Å, and 4.146 Å, respectively (Figure 2). These short contact distances indicate that the identified residues are positioned in close spatial proximity at the predicted ADAR1–PKR interaction interface, supporting their potential contribution to complex formation. The top-ranked docking conformation yielded a docking score of –180 and a confidence score of 0.6. In contrast, no direct contacts involving Thr446 were identified in our docking model. These findings suggest an alternative binding orientation in which ADAR1 engages PKR through a region close but distinct from that predicted by Sinigaglia [19]. Although this model requires experimental validation, in agreement with the Singalgia et al study it supports the possibility that ADAR1 can directly interact with PKR through multiple, context-dependent binding modes independent of its RNA-editing activity (Figure 2). Moreover, to evaluate whether dsRNA binding is compatible with the predicted ADAR1–PKR interaction, the top-ranked ADAR1–PKR docking model was subsequently used for dsRNA docking. The resulting ternary complex showed that dsRNA remained associated with the ADAR1 dsRBDs while preserving the predicted ADAR1–PKR interface, suggesting that the interaction is structurally compatible with the RNA-bound state of ADAR1.

**Figure 2:**
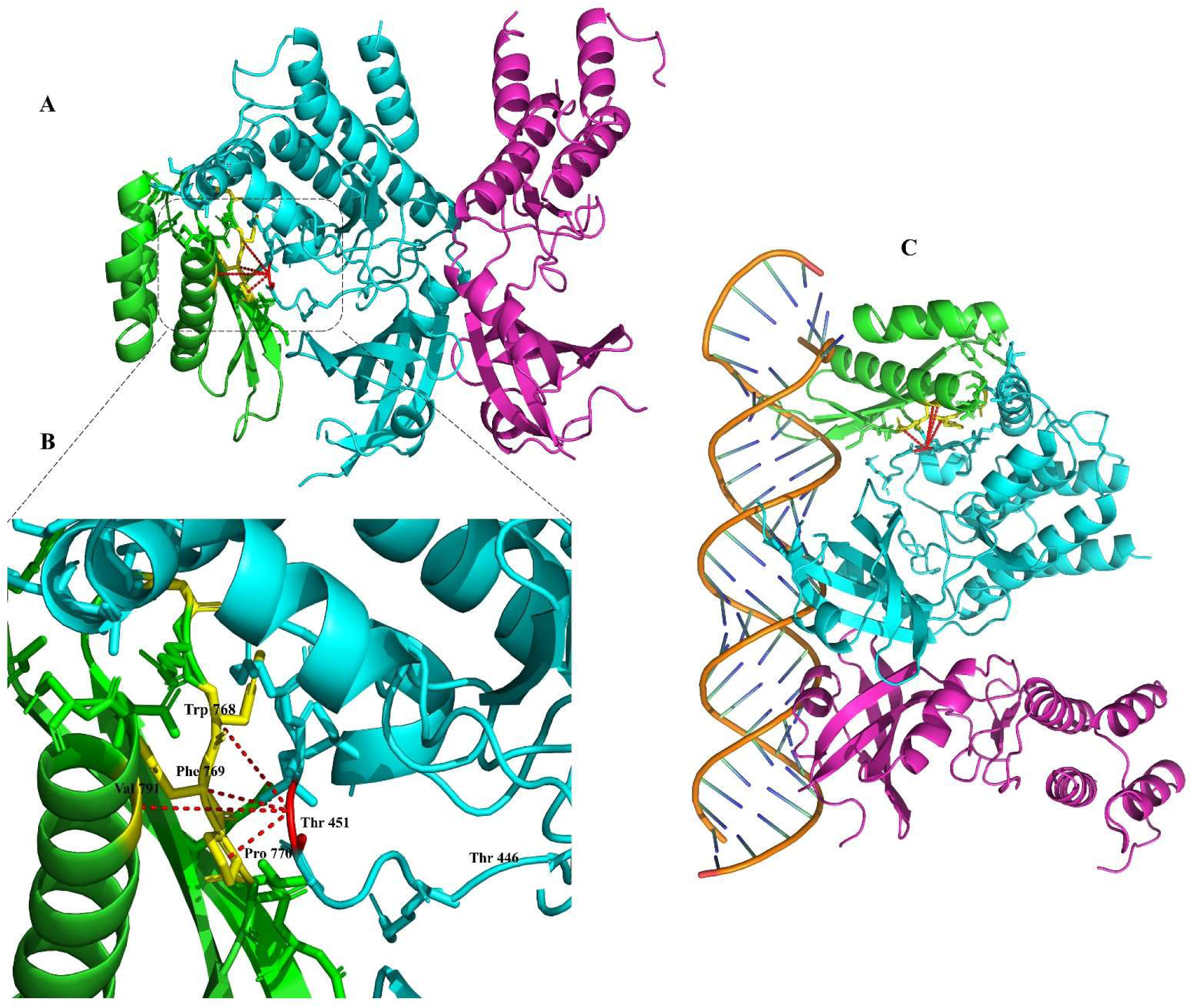
Predicted interaction between the PKR kinase domain and the RBDIII domain of ADAR1. (A) Overall docked complex showing PKR Chain B (cyan) and Chain C (magenta) interacting with the RBDIII domain of ADAR1 (green). PKR residue Thr451 is highlighted in red, while the predicted interface residues of ADAR1 are shown in yellow. Dashed lines indicate the predicted residue–residue contacts at the protein–protein interface. (B) Enlarged view of the predicted interaction interface between PKR and the ADAR1 RBDIII domain, highlighting the key contact residues. (C) Docked ADAR1–PKR complex in the presence of the dsRNA molecule (PDB ID: 6E0F), illustrating the spatial arrangement of the predicted protein–protein interaction relative to the bound RNA.

### The ADAR1-PKR interaction surface is conserved in ADAR2 and ADAR3 and ADAR1 orthologs in vertebrates

The cytoplasmic ADAR1p150 isoform is considered the principal regulator of PKR through an RNA editing-independent mechanism [24]. In contrast, ADAR1p110, ADAR2, and ADAR3 are predominantly localized to the nucleus, limiting their access to cytoplasmic innate immune effectors such as PKR [27]. Nevertheless, it has been shown that cytoplasmic ADAR2 (cytoADAR2), an engineered ADAR2 variant lacking the nuclear localization signal, suppresses PKR phosphorylation and stress granule formation following interferon treatment [19, 24]. These findings can suggest that the PKR-interacting interface may be conserved between ADAR1 and ADAR2, although direct structural evidence supporting this hypothesis is currently lacking. Consistent with this possibility, primary sequence alignment revealed several highly conserved regions across the human ADAR family, suggesting the preservation of residues important for structural integrity and protein function (Figure 3). It is important to note, however, that the region predicted to mediate PKR binding exhibited partial sequence conservation. Specifically, the motif GRWFPA (residues 766–771) within the ADAR1 dsRBDIII domain corresponds to GQFFEG in ADAR2 and GRTFEG in ADAR3, with two and three amino acid substitutions, respectively.

**Figure 3:**
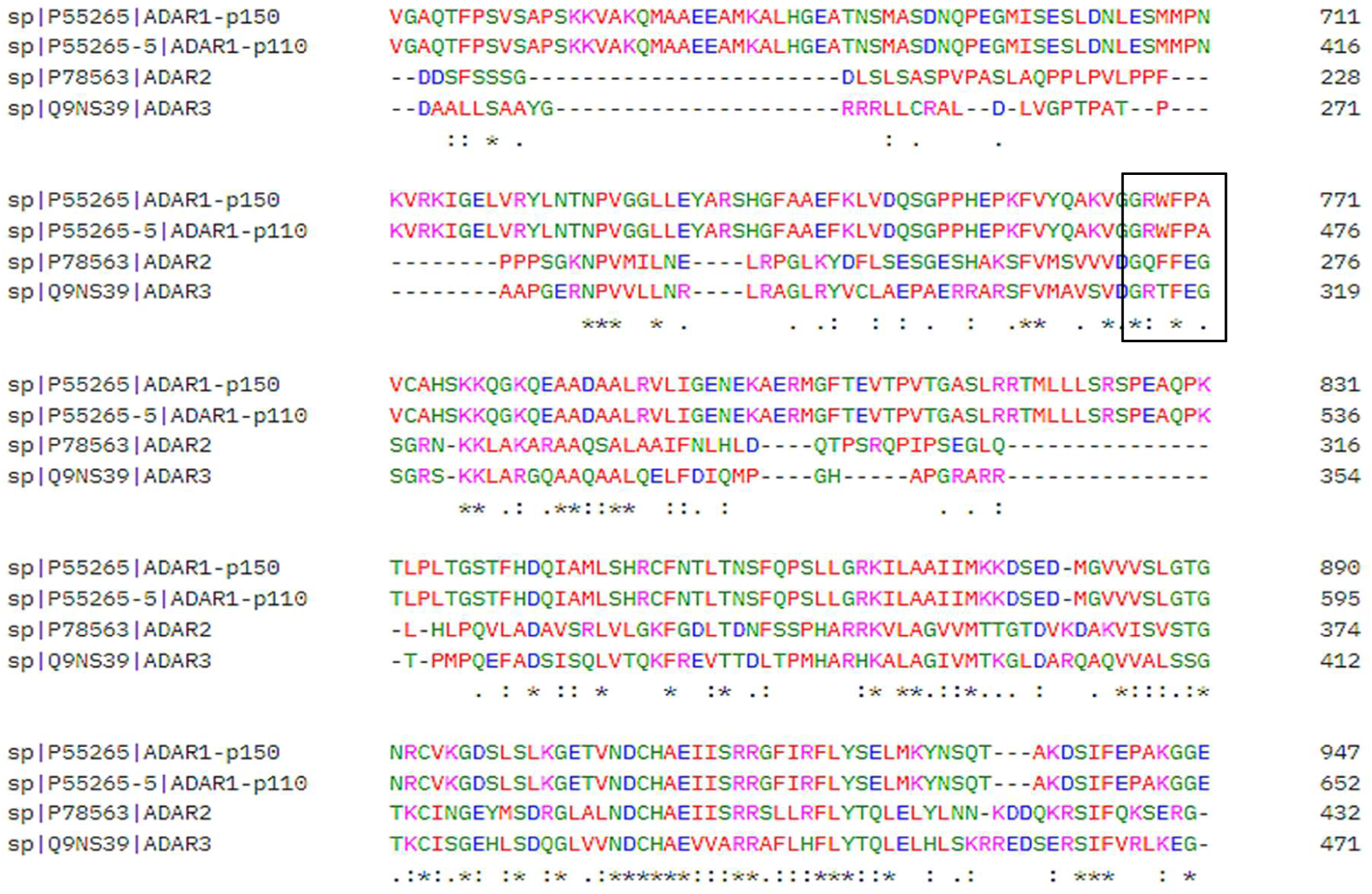
Multiple sequence alignment (MSA) of the ADAR protein family ((ADAR1_Human: P55265, (ADAR2_Human: Q9BZV3), (ADAR3_Human: Q9NQZ3), performed using Clustal Omega. The alignment highlights conserved regions across different ADAR homologs, emphasizing sequence conservation within functional domains. In the alignment, an asterisk (*) indicates fully conserved residues, a colon (:) denotes conservation between residues with strongly similar properties, and a period (.) represents conservation between residues with weakly similar properties.

Despite these sequence differences, some substitutions preserve similar physicochemical properties. For example, tryptophan (W) and phenylalanine (F) are both aromatic and hydrophobic residues, while alanine (A) and glycine (G) are both small residues, although glycine has greater conformational flexibility. These similarities suggest partial conservation of physicochemical features despite differences in the exact amino acid sequences (Figure 3). Furthermore, the predicted PKR-interaction motif, GRWFPA, is highly conserved among vertebrate ADAR1 orthologs, including homo sapiens (human), *P. troglodytes* (chimpanzee), *M. musculus* (mouse), *G. gallus* (chicken), and *D. rerio* (zebrafish) (Figure 4). The high degree of conservation across vertebrate species may indicate that this region is under strong evolutionary constraint and could play an important functional role.

**Figure 4:**
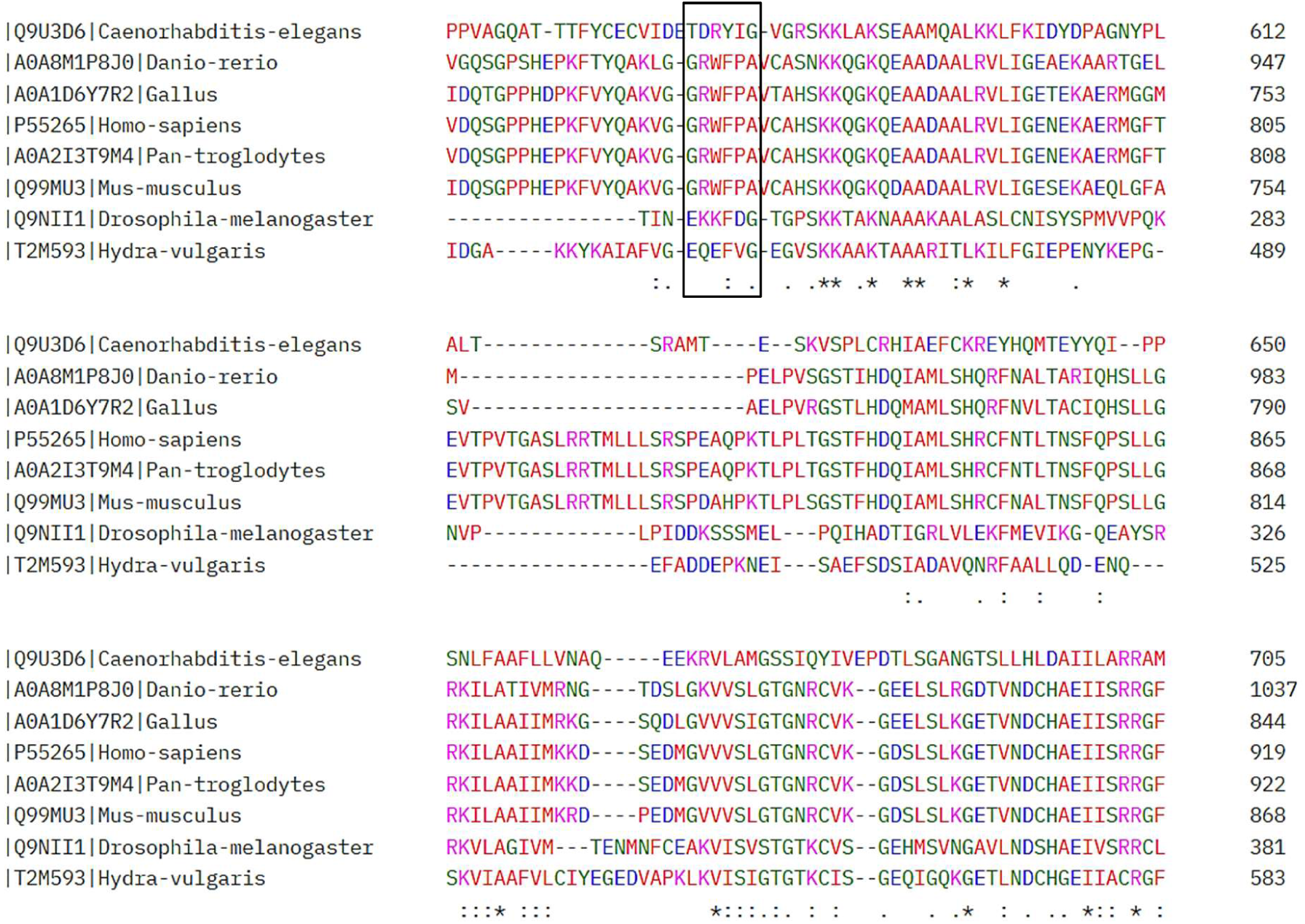
Multiple sequence alignment (MSA) of ADAR proteins from *H. sapiens* (P55265), *P. troglodytes* (A0A2I3T9M4), *M. musculus* (Q99MU3), *G. gallus* (A0A1D6Y7R2), *D. rerio* (A0A8M1P8J0), *C. elegans* (Q9U3D6), *D. melanogaster* (Q9NII1), and *H. vulgaris* (T2M593), generated using Clustal Omega. The alignment highlights conserved regions across ADAR homologs, emphasizing sequence conservation within functional domains. The boxed region indicates the human ADAR1 GRWFPA motif containing the predicted PKR-interacting residues, which is absent or highly divergent in PKR-negative species. In the alignment, an asterisk (*) denotes fully conserved residues, a colon (:) indicates residues with strongly similar physicochemical properties, and a period (.) represents residues with weakly similar properties.

Based on this observation, we hypothesized that conservation of the predicted PKR- interaction interface may be linked to the evolutionary emergence of PKR. Consistent with this hypothesis, the corresponding motif was poorly conserved in species lacking a PKR ortholog. Specifically, the corresponding regions in D. melanogaster (fruit fly), C. elegans (nematode), and H. vulgaris (hydra) showed little or no sequence similarity to the conserved vertebrate motif (Figure 4). Notably, residues predicted by protein-protein docking to participate in the ADAR1- PKR interface (Trp768, Phe769, and Pro770) were either absent or not conserved in these PKR- negative species. In particular, the aromatic residues Trp768 and Phe769, which contribute to the predicted interaction interface, were completely absent, disrupting the characteristic GRWFPA core motif. As the docking analysis predicted that PKR binding is mediated by a cluster of neighboring residues rather than a single amino acid, the loss of this conserved aromatic-rich motif is consistent with a diminished capacity to support the predicted ADAR1-PKR interaction. Collectively, these observations are consistent with the hypothesis that the ADAR1-PKR interaction interface has co-evolved with PKR during vertebrate evolution. However, this evolutionary relationship remains speculative and will require functional validation and broader comparative analyses across diverse taxa.

### 3.3 Docking analysis of ADARs family

Primary sequence comparison of ADAR family members revealed only minor amino acid differences within the predicted PKR-interaction interface. Although previous studies have suggested that ADAR family members may share the ability to regulate PKR activity [19, 35], their distinct subcellular localizations may influence this function. In particular, ADAR1p150 is predominantly cytoplasmic, where it can readily interact with PKR, whereas ADAR1p110, ADAR2 and ADAR3 are primarily nuclear, potentially limiting their access to cytoplasmic PKR under physiological conditions. Moreover, our structural analysis suggests that the predicted PKR-binding interface is not highly conserved across human ADAR proteins. To evaluate whether the sequence conservation and function translates into a conserved structural mode of PKR binding, we performed comparative protein-protein docking analyses of PKR with ADAR1, ADAR2, and ADAR3.

Protein-protein docking analysis further demonstrated that ADAR2 forms a less extensive predicted interaction interface with PKR than ADAR1. In the predicted ADAR2-PKR complex, PKR Thr451 interacted with only two residues, Arg244 and Asp270 showing distance of 4.264 Å and 4.770Å respectively, both located within the second dsRNA-binding domain (dsRBDII), whereas in the ADAR1 complex it interacted with four residues (Trp768, Phe769, Pro770, and Val791). Of these, only Asp270 corresponded to a conserved residue identified by multiple sequence alignment, whereas Arg244 was located outside the conserved motif (Figure 5). The docking score (–113.10), confidence score (0.643), were all less favorable than those obtained for ADAR1 (Table 1), consistent with lower confidence in the predicted ADAR2-PKR complex.

**Figure 5:**
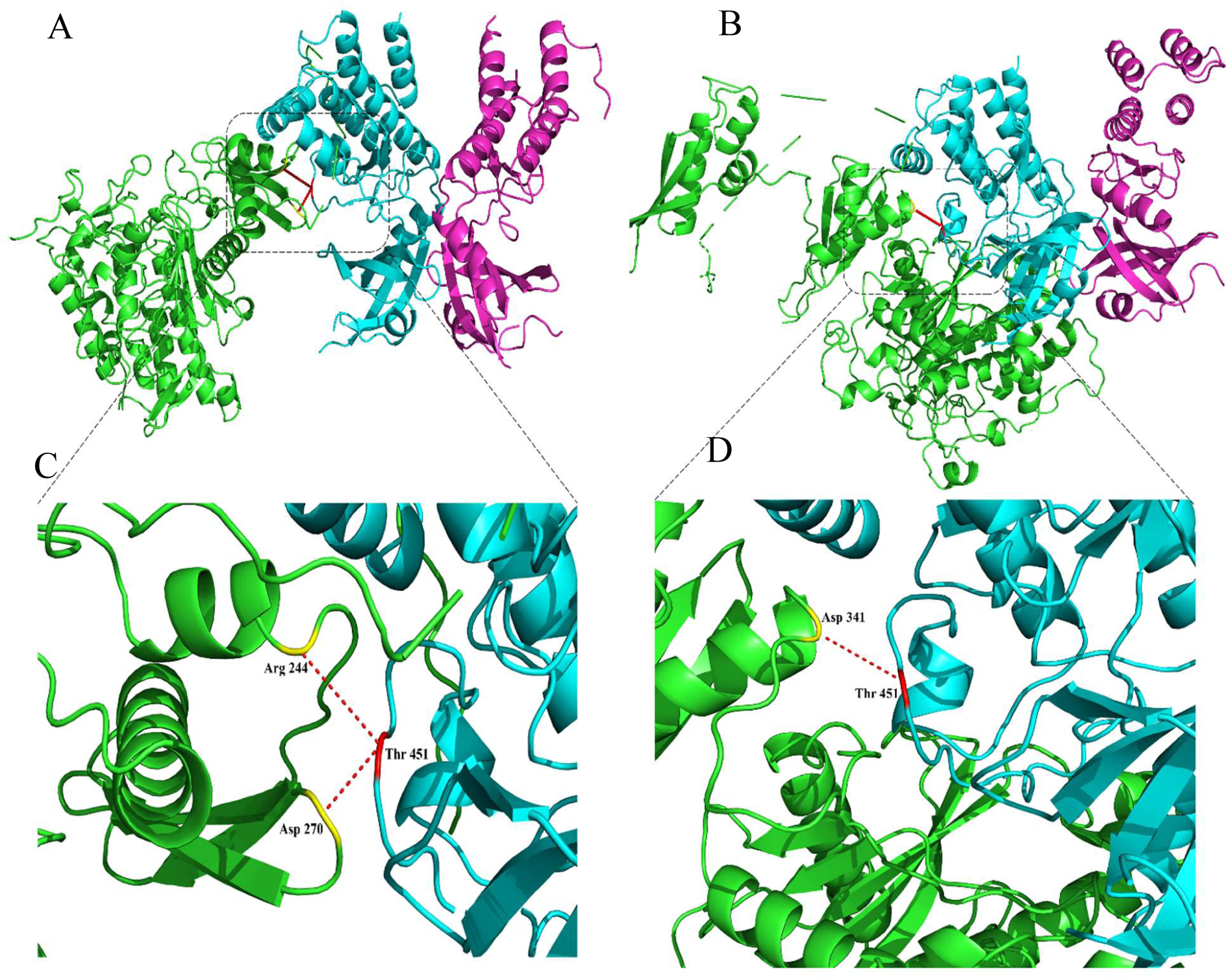
Predicted interactions between the PKR kinase domain and the RBDII domains of ADAR2 and ADAR3. (A, B) Overall docked complexes showing PKR Chain B (cyan) and Chain C (magenta) interacting with the RBDII domain of ADAR2 (A) and ADAR3 (B) (green). PKR residue Thr451 is highlighted in red, while the predicted interface residues of ADAR2 and ADAR3 are shown in yellow. Dashed lines indicate the predicted residue–residue interactions at the protein–protein interface. (C, D) Enlarged views of the interaction interfaces between PKR and the RBDII domains of ADAR2 (C) and ADAR3 (D), highlighting the key contact residues involved in the predicted interactions.

**Table 1.** Predicted interactions of PKR Thr451 with ADAR proteins.

| **Protein**<br>**(PKR interaction domain)** | **Residues in contact with**<br>**PKR Thr451** | **Docking**<br>**score** | **Confidence**<br>**score** |
| --- | --- | --- | --- |
| **ADAR1 (GRWFPA)** | Trp768, Phe769, Pro770, Val791 | -180 | 0.6 |
| **ADAR2** | Arg244, Asp270 | -113.10 | 0.643 |
| **ADAR3** | Asp341 | -105.74 | 0.547 |
| **Mutated ADAR1**<br>**PKR in contact with Phe489** | Tyr769, Gly770 | -167.10 | 0.5 |

ADAR3 exhibited an even less extensive predicted interaction with PKR. Docking analysis identified a single contact between PKR Thr451 and Asp341 with distance of 4.163Å, located within the dsRNAII-binding domain. Unlike the interacting residues identified in ADAR1 and ADAR2, Asp341 was not part of the conserved motif revealed by the MSA (Figure 4). ADAR3 also produced the least favorable docking metrics among the three ADAR proteins, with a docking score of –105.74 and a confidence score of 0.547. Collectively, these findings indicate that the predicted PKR-binding interface is not structurally conserved among human ADAR family members and that ADAR3 exhibits the lowest predicted structural compatibility for PKR binding. However, these predictions require validation by biochemical and functional studies to establish their biological relevance.

### Mutations Within the Predicted Interaction Interface Alter ADAR1–PKR Binding

In our study, the conserved ADAR1 motif GRWFPA was identified as the interaction interface between ADAR1 and PKR. To investigate the functional significance of the conserved motif GRWFPA within the RBDIII domain of ADAR1, site-directed mutagenesis was performed using the PyMOL Mutagenesis Wizard, followed by protein-protein docking analysis. Briefly, three residues within the motif were substituted to generate the sequence GR<u>EYG</u>A, these substitutions were selected to alter the physicochemical properties of residues predicted to contribute to the ADAR1-PKR interaction interface while maintaining the overall protein backbone. In first mutate W768E, in which tryptophan (W), a bulky hydrophobic aromatic residue, was replaced with glutamic acid (E), a negatively charged residue, thereby disrupting hydrophobic packing and introducing electrostatic repulsion at the predicted PKR-binding interface [36]. F769 Y, phenylalanine was change into tyrosine, because both residues are aromatic and structurally similar, allowing evaluation of whether introduction of a single hydroxyl group and a modest increase in polarity is sufficient to perturb the predicted PKR-binding interface while largely preserving the overall fold of the dsRBDIII domain. Thus, F769Y represents a conservative mutation designed to assess the contribution of hydrophobic packing rather than complete disruption of the interface [37]. In P770G mutant, proline 770 was mutated to glycine to eliminate the conformational rigidity imposed by proline and increase local backbone flexibility, thereby testing whether structural rigidity at this position contributes to PKR recognition. The mutated residues were chosen to disrupt the potential hydrogen bonding and hydrophobic interactions while preserving the motif’s backbone structure. The structure of mutated proteins was visually inspected and not changes in structural integrity were observed, and the modified ADAR1 RBDIII domain was saved in PDB format for successive analysis.

The mutant ADAR1 RBDIII structure was subsequently subjected to protein-protein docking with the PKR kinase domain using the HDOCK server under default parameters. The resulting docking models were analyzed to identify changes in binding patterns, with particular emphasis on the loss or gain of intermolecular interactions relative to the wild-type ADAR1–PKR complex. Structural comparison revealed the loss of the interaction with PKR residue Thr451 and the formation of new contacts between ADAR1 residues Tyr769 (distance 4.761 Å) and Gly770 (distance 4.891 Å) with PKR residue Phe489 (Figure 6). These changes indicate that the introduced mutations altered the binding interface, suggesting a reorganization of the interaction network that may affect the stability and/or affinity of the ADAR1-PKR complex. After mutating the GRWFPA motif of ADAR1’s RBDIII domain to a new sequence (GR<u>EYG</u>A), we studied how these changes affected the interaction between ADAR1 and PKR using the HDOCK server. The new contact with Phe489 proposes that the proteins now bind in a different way with a docking score -167.10, confidence score 0.5.

**Figure 6:**
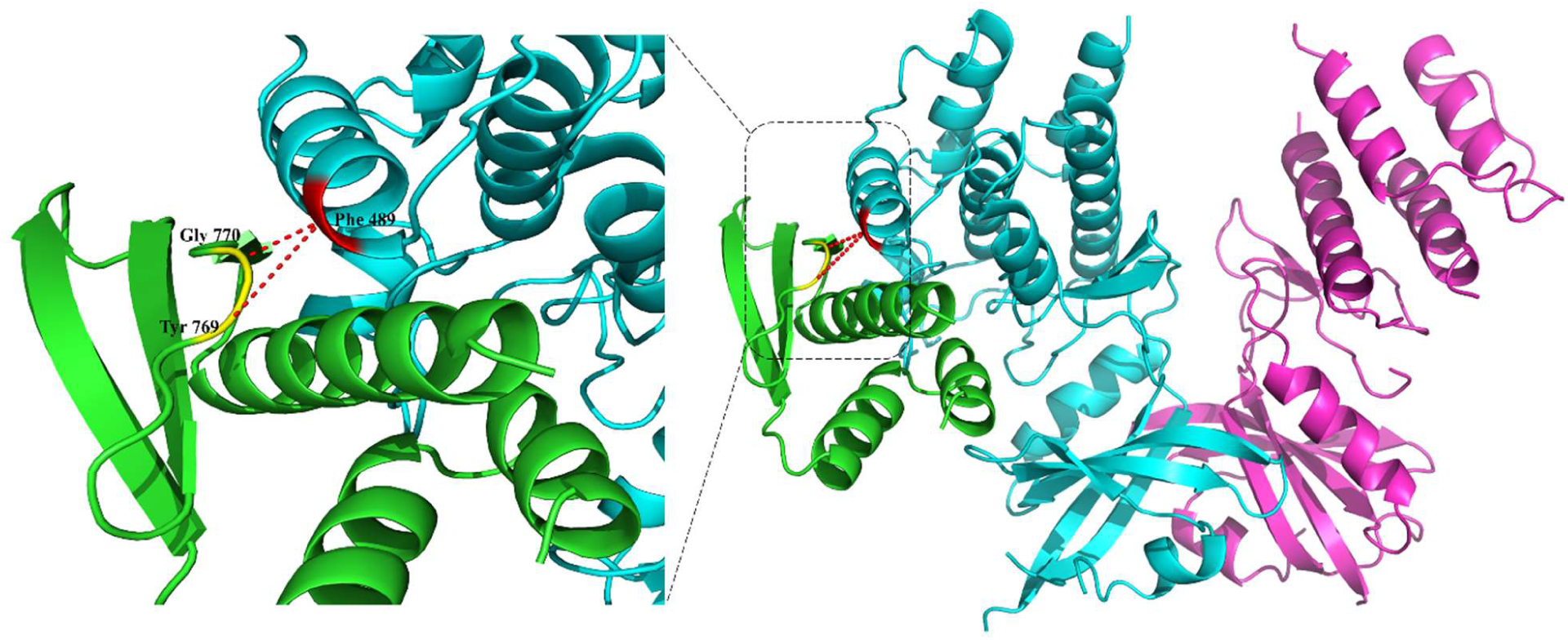
Structural model of the ADAR1 triple mutant (W768E/F769Y/P770G) docked with the PKR kinase domain. (A) Overall docked complex showing PKR Chain B (cyan) and Chain C (magenta) interacting with the mutated ADAR1 RBDIII domain (green). Interface residues are highlighted in yellow, and Phe489 of PKR is shown in red. (B) Zoomed view of the predicted interaction interface highlighting the altered binding mode following mutation.

These results show that the original residues in the GRWFPA motif are important for proper binding and function, and that changing them affects how ADAR1 and PKR interact. Although Phe489 is not part of the phosphorylation sites essential for PKR activation (such as Thr451 and Thr446) [18, 20], its involvement in the interaction with mutated ADAR1 residues suggests a shift in the binding interface. This interaction is likely non-functional, as Phe489 does not play a known role in in the activation loop, may have had more direct implications for PKR’s regulatory activity. PKR activation or substrate recognition. In contrast, the original interaction with Thr451, located within activation loop, have more direct implications for PKR regulatory activity and its interaction with ADAR1.

### ADAR1p150-ΔWFP Shows Reduced Co-immunoprecipitation with PKR

To assess the contribution of the predicted interaction interface to ADAR1–PKR association, we compared the ability of wild-type and mutant ADAR1 proteins to associate with endogenous PKR by co-immunoprecipitation. We hypothesized that disruption of the conserved interface would weaken ADAR1–PKR interaction. Briefly, ADAR1-knockout cells [28, 31] were transfected with plasmids expressing GFP-tagged wild-type (WT) ADAR1p150 or a series of mutants (W768E, F769Y, P770G, K778E, R790E, and the combined ΔWFP mutant). W768E, F769Y, P770G, and ΔWFP targeted residues within our predicted interaction interface, whereas K778E and R790E were selected based on the study by Sinigaglia et al [19]. In that study, of these two mutants, only R790E failed to inhibit PKR autophosphorylation and showed impaired co-immunoprecipitation with PKR. GFP-ADAR1 proteins were immunoprecipitated using an anti-GFP antibody, and co-immunoprecipitated endogenous PKR was analyzed in the presence or absence of RNase treatment. All constructs were expressed at comparable levels, exhibited predominantly cytoplasmic localization, and were recovered with similar efficiency during immunoprecipitation, indicating that the introduced substitutions did not measurably affect protein expression, stability, subcellular localization, or immunoprecipitation efficiency (Figure 7). RNase treatment reduced the amount of co-immunoprecipitated PKR by approximately tenfold but did not abolish the interaction, indicating that RNA substantially contributes to, but is not solely responsible for, ADAR1–PKR complex formation, consistent with previous studies. Nonetheless, the transient overexpression system did not provide sufficient resolution to reproducibly detect differences in PKR binding between the individual single-amino-acid ADAR1 mutants and the wild-type protein (Figure 7). This observation is broadly consistent with our docking analysis, which predicted only a modest reduction in the docking score following the combined mutation rather than complete disruption of the ADAR1–PKR interaction. However, our findings contrast with those of Sinigaglia et al., who reported that the R790E mutant failed to co-immunoprecipitate with PKR. At present, we cannot fully explain this discrepancy, although it may arise from differences in the experimental models used. Notably, only the combined mutant (ΔWFP) consistently showed a trend toward reduced PKR co-immunoprecipitation compared with wild-type ADAR1, although the difference did not reach statistical significance (p=0,66; representative Figure 7). A similar trend was observed following RNase treatment (p=0,51representative Figure 7). The direction of this effect is consistent with the lower docking score predicted for the combined mutant, suggesting that the conserved motif may contribute cooperatively to PKR association. However, the magnitude of the effect observed in this experimental system was insufficient to definitively establish the contribution of this region, indicating that additional biochemical or quantitative binding approaches will be required to determine its precise role in ADAR1–PKR interaction.

**Figure 7.**
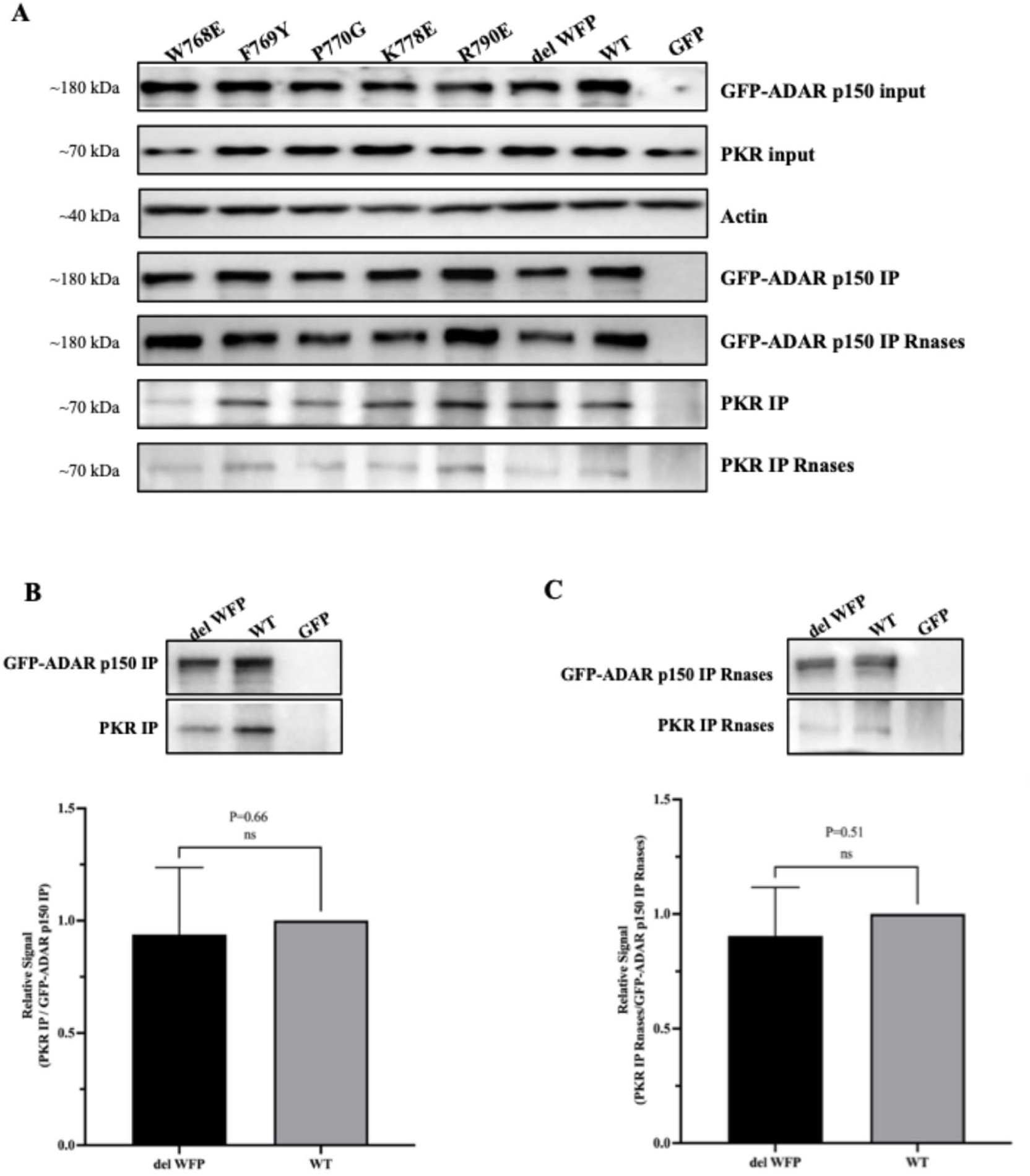
ADAR1p150-ΔWFP Shows Reduced Co-immunoprecipitation with PKR. **(A)** Expression of GFP-ADAR1 p150 variants with endogenous PKR. ADAR1-knockout HEK-293 cells were transfected with plasmids expressing the indicated mutant constructs (W768E, F769Y, P770G, K778E, R790E, and del WFP), wild-type (WT) GFP-ADAR1 p150, or GFP control. Cell lysates were immunoprecipitated using an anti-GFP antibody in the presence or absence of RNase treatment analyzed by Western blot. **(B)** Representative Western blot (*top*) and relative quantification (*bottom*) of endogenous PKR co-immunoprecipitated with WT GFP-ADAR1 p150 or the del WFP mutant under untreated conditions. The PKR Co-IP signal was normalized to WT (*P* = 0.66, ns). **(C)** Representative Western blot (*top*) and relative quantification (*bottom*) of endogenous PKR co-immunoprecipitated with WT GFP-ADAR1 p150 or the del WFP mutant following RNase treatment. The PKR Co-IP signal was normalized to WT (*P* = 0.51, ns).

Collectively, these results indicate that disruption of the predicted interface does not produce a large loss of ADAR1–PKR association under the conditions tested. While the observed trend for the combined mutant is consistent with the docking predictions, the contribution of this conserved motif to PKR binding appears to be modest and remains to be established using more quantitative approaches.

## Discussion

The dsRNA sensor PKR, although best known for its antiviral activity, plays important roles in a broad range of cellular processes, including the regulation of mRNA translation, transcription, apoptosis, and cell proliferation [8, 38, 39]. Consequently, inappropriate or sustained PKR activation can disrupt cellular homeostasis and has been implicated in several degenerative and inflammatory diseases [40]. ADAR1, on the other hand, is a key dsRNA-binding protein that catalyzes A-to-I RNA editing and plays a central role in preventing inappropriate recognition of endogenous dsRNA and aberrant activation of dsRNA-sensing pathways, including [7, 41]. PKR. Surprisingly, despite the importance of ADAR1-mediated regulation of PKR in efficient viral replication and its potential contribution to cancer development and progression, the exact molecular mechanisms by which ADAR1 controls PKR activation remain poorly defined. Previous studies indicate that ADAR1 can regulate PKR through both competition for dsRNA substrates and association with PKR itself [19, 24]. The enrichment of A-to-I-edited transcripts among PKR-bound RNAs supports competition between ADAR1 and PKR for shared dsRNA substrates [42, 43]. In parallel, multiple studies have demonstrated an association between ADAR1 and PKR, with evidence for both RNA-dependent and RNA-independent components [19, 28, 33, 43–45]. Both ADAR1 isoforms, p110 and p150, have been detected in association with PKR, although preferential association of the cytoplasmic p150 isoform has also been reported [28, 33, 43, 44]. The differences among these observations may reflect differences in cell type, experimental conditions, antibodies, or expression systems. Nevertheless, the cytoplasmic localization and interferon inducibility of ADAR1p150 make this isoform particularly relevant to PKR regulation during viral infection.

Evidence from deletion and functional studies further implicates the ADAR1 dsRNA-binding domains in PKR regulation. The N-terminal region unique to ADAR1p150, including the Zα domain, does not appear to be essential for PKR inhibition, whereas deletion of individual or multiple dsRBDs compromises this activity [33, 43, 44]. Similarly, inactivation or deletion of the C-terminal deaminase domain does not abolish PKR inhibition, indicating that RNA-editing activity itself is not sufficient to explain this function [33]. These observations collectively support a role for the dsRBD-containing region in ADAR1–PKR regulation, potentially through a combination of RNA-dependent recruitment and direct protein–protein interactions.

In the present study, docking analysis identified a favorable interaction interface between ADAR1 RBDIII and PKR in which the conserved ADAR1 GRWFPA motif was positioned within the predicted contact region. Specifically, W768, F769, and P770 were located close to PKR Thr451, suggesting that this region may contribute to ADAR1-PKR recognition (Figure 2). Importantly, the docking results did not indicate that these residues alone were sufficient to maintain the complete interaction. Instead, combined disruption of the WFP motif altered the predicted interaction pattern, with the prominent PKR contact shifting from Thr451 toward Phe489 (Figure 6). Thus, the computational analysis suggested that the WFP motif may contribute to the organization and stability of the interface rather than function as an isolated interaction determinant.

Our findings are particularly relevant in comparison with the study of Sinigaglia et al [19], which provided strong evidence for a direct interaction between ADAR1 dsRBD3 and the PKR kinase domain. Their structural analysis identified an interaction involving the C-lobe of PKR and highlighted ADAR1 residues including R790 and K778. Functionally, R790E substantially impaired ADAR1-PKR association and reduced ADAR1-mediated inhibition of PKR activation. Our study differs by independently identifying the neighboring W768/F769/P770 motif as a potential component of the interaction interface and experimentally testing these residues. The weaker effect of WFP mutations compared with the previously reported R790E phenotype suggests that the ADAR1-PKR interface contains residues with different active contributions. R790 may represent a stronger determinant of a specific protein-protein contact, whereas the WFP motif may contribute collectively to overall complex stability. Rather than contradicting the previous study, our findings therefore extend its model by suggesting that the interface may be more extensive and functionally heterogeneous than a single dominant residue or contact.

Comparison with ADAR2 and ADAR3 further supports the possibility of ADAR1-specific structural determinants within the PKR interaction interface. ADAR2 and ADAR3 produced different predicted interaction surfaces and weaker docking scores with PKR, with residues such as Arg244 and Asp270 for ADAR2 and Asp341 for ADAR3 (Table 1; Figure 5). Although docking scores cannot be interpreted as direct measurements of binding affinity, the distinct predicted interfaces are consistent with differences in the molecular architecture of the three ADARs proteins. This distinction is particularly relevant to ADAR1p150, which is the cytoplasmic, interferon-inducible isoform with an established role in regulating dsRNA sensing and PKR activity. Furthermore, a recent study demonstrated that ADAR1p150 suppresses PKR activation, at least in part, by competing with PKR for binding to dsRNA substrates [24]. This model also explains the ability of cyto-ADAR2 to inhibit PKR despite lacking the p150-specific N-terminal region [24]. The experimental Co-IP analysis provided a direct test of the computational prediction. ADAR1-deficient cells expressing GFP-tagged ADAR1p150 WT or W768E, F769Y, P770G, K778E, R790E and ΔWFP mutants were used to examine association with endogenous PKR. The individual W768E, F769Y and P770G substitutions did not reproducibly produce a clear reduction in PKR association. In contrast, the combined ΔWFP mutant consistently showed a trend toward reduced PKR co-immunoprecipitation, although this difference was not statistically significant. These findings indicate that the W768/F769/P770 motif is not individually essential for ADAR1–PKR association. Nevertheless, the consistent direction of the ΔWFP phenotype, together with the altered docking configuration following WFP disruption, supports a possible cooperative contribution of this motif to the interaction interface. The experimental results therefore favor a model in which the WFP region contributes to, but does not independently determine, PKR association.

An important finding was the strong effect of RNase treatment. RNase reduced co-immunoprecipitated PKR by approximately tenfold but did not completely abolish the interaction. This demonstrates that RNA substantially contributes to ADAR1–PKR complex formation while a measurable RNA-independent component remains. This observation is consistent with previous studies showing that ADAR1-PKR association depends strongly on dsRNA binding but also involves direct protein–protein contacts [19, 28, 33]. Thus, our findings support a model in which RNA facilitates the recruitment or stabilization of ADAR1 and PKR, while direct contacts within the ADAR1-PKR interface provide additional stabilization (Figure 7).

Several limitations should be considered when interpreting these findings. Docking analysis predicts possible interaction geometries but does not directly measure binding affinity or establish the energetic contribution of individual residues. Similarly, Co-IP detects recovery of a cellular protein complex but cannot, by itself, distinguish direct protein–protein binding from RNA-mediated or indirect association. Most importantly, the non-significant phenotype of the ΔWFP mutant does not establish the W768/F769/P770 motif as an essential determinant of PKR binding. The present data therefore support a contributory rather than indispensable role for this region. Quantitative biochemical approaches, such as surface plasmon resonance, biolayer interferometry, microscale thermophoresis, or fluorescence polarization, together with targeted structural or mutational analyses, will be required to determine the precise energetic contribution of the WFP motif and its individual residues.

In conclusion, our findings support a model in which ADAR1p150-PKR association involves both RNA-mediated interactions and direct protein-protein contacts. The W768/F769/P770 motif may contribute cooperatively to interface organization, although its role in PKR association requires further validation. These results extend the existing model of ADAR1 RBDIII-PKR interaction and provide a basis for further investigation of the molecular mechanisms underlying PKR regulation during antiviral responses.

## Acknowledgements

The study was supported by the Croatian Science Foundation grant (HRZZ) IP-2025-02-3026 and DOK-NPOO-2023-10-7304, and University of Rijeka Research grant uniri-mzi-25-43 to I.J.

